# A toxic palmitoylation on Cdc42 drives a severe autoinflammatory syndrome

**DOI:** 10.1101/808782

**Authors:** Bahia Bekhouche, Aurore Tourville, Yamini Ravichandran, Rachida Tacine, Laurence Abrami, Michael Dussiot, Andrea Khau-Dancasius, Olivia Boccara, Meriem Khirat, Marianne Mangeney, Nathalia Bellon, Sylvie Fraitag, Smail Hadj-Rabia, Stéphane Blanche, Anne Puel, Sandrine Etienne-Manneville, F. Gisou van der Goot, Jacqueline Cherfils, Olivier Hermine, Jean-Laurent Casanova, Christine Bodemer, Asma Smahi, Jérôme Delon

## Abstract

**Background:** Autoinflammatory diseases (AID) result from dysregulation of the first lines of innate immune responses. Recently, development of high throughput genome sequencing technology led to the rapid emergence of important knowledge in the genetic field. About 20 genes have been identified so far in monogenic forms of distinct AID. However, 70-90 % of patients with AID remain without genetic diagnosis.

**Objective:** We report the identification and characterization of a mutation in the C-terminal region of the Rho GTPase Cdc42 in a patient presenting a severe autoinflammatory phenotype.

**Methods:** We have analyzed the consequences of the mutation on the subcellular localization of the Cdc42 protein using imaging techniques. Molecular studies were performed using proteomic and biochemical experiments to provide mechanistic bases of the observed defects. Functional assays were also conducted using flow cytometry and cytokine production measurements.

**Results:** We show that mutant Cdc42 is trapped in the Golgi apparatus due to the aberrant addition of a palmitate that both enhances the interaction of mutant Cdc42 with Golgi membranes and inhibit its extraction by GDP dissociation inhibitor (GDI), thus impairing its cytosol/membrane shuttling. At the functional level, mutant Cdc42 fails to sustain actin filaments polymerization and induces an exacerbated profile of pro-inflammatory cytokine production due to increased NF-κB activation.

**Conclusions:** Our study now provides a molecular explanation for mutations that have been identified recently in our AID patient and others in the C-terminal part of Cdc42. Mutations located in this region of Cdc42 impair the intracellular localization of Cdc42, preventing its interaction with the plasma membrane. Thus, our results definitively link mutations in the *CDC42* gene to a complex immune-hemato-autoinflammatory phenotype in humans.

## Introduction

The delineation of familial Mediterranean fever (FMF) and the identification of responsible mutations in *MEFV* (encoding human pyrin) in 1997 ^1,2^ led to the emergence of autoinflammatory diseases (AID) as a distinct nosology entity. This group of diseases became clearly distinct from autoimmune diseases and result from dysregulation of first lines of innate immune responses ^3^. Interestingly, in the large majority of cases, IL-1β, a master cytokine in inflammation, is overproduced. This strong feature makes the use of IL-1β inhibitors such as anakinra successful. Recently, development of high throughput genome sequencing technology led to the rapid emergence of important knowledge in the genetic field. About 20 genes have been identified so far in monogenic forms of distinct autoinflammatory diseases ^4^. The association between genetic defects and pathophysiological mechanisms was evident in most cases. Now, at least four groups of diseases have been defined, mainly related to 4 different pathways: (i) IL-1β- (ii) IFNα-, (iii) NF-κB signaling- and (iv) persistent macrophage activation-mediated AID.

Despite the discovery of some responsible genes, 70-90 % of patients with AID remain without genetic diagnosis. Some entities remain unclassified and more investigations are needed to decipher the relationships between genetic defects and clinical manifestations. Similarities in clinical phenotypes strongly suggest cross talks between common canonical signaling pathways.

Small GTPases of the Rho family have recently been increasingly associated to inflammatory diseases and complex phenotypes in humans ^5–8^, and thus represent potential candidates in such signaling dysfunctions. Previous findings classify psoriasiform manifestations in the group of AID. In such cases, elevated levels of active RhoA and Rac1 have been reported ^9,10^. Recently, Gernez *et al* reported 4 patients with severe autoinflammation that were successfully treated with IL-1β inhibition and identified 3 distinct *de novo* heterozygous variants in the C-terminal part of the Cdc42 ^11^.

Rho GTPases regulate myriads of cellular functions, through their activation by upstream signals transmitted by multiple cell surface receptors and their ability to activate various downstream signaling pathways, often involving the regulation of actin cytoskeleton dynamics. For example, active Cdc42 recruits the Wiskott-Aldrich syndrome protein (WASP) effector which, in turn, stimulates the Arp2/3 complex to induce the polymerization of actin filaments, resulting in the formation of membrane protrusions such as filopods ^12,13^. The ubiquitous and brain-specific isoforms of Cdc42 have a central role in the development and maintenance of cellular polarity as well as in directional migration ^14^. Polarity is described as the asymmetrical organization of cellular components along different axes. This phenomenon is present in many cell types and is essential for development and cellular communication.

As all members of this superfamily, Rho family, of which RhoA, Rac1 and Cdc42 are the best studied representatives, alternate between an inactive form bound to GDP and an active form bound to GTP. This alternation is regulated by stimulation of GDP/GTP exchange by guanine nucleotide exchange factors (GEFs) whereas inactivation occurs by GTP hydrolysis stimulated by GTPase-activating proteins (GAPs) (reviewed in ^15,16^). Binding of GTP allows small GTPases to interact with, and activate proteins called « effectors » (reviewed in ^15–17^). The cytosol/membrane cycle is regulated by GDP Dissociation Inhibitors (GDIs), which extract them from endo and plasma membranes, and prevent their activation by sequestering them in the cytosol. Membrane association of Rho GTPases requires the post-translational addition of lipids by geranyl-geranyl transferase (GGTase) I (reviewed in ^18–20^), a process called prenylation that occurs on cysteine residues located in C-terminal CAAX sequences ^19^. Importantly, most, if not all, GEFs, GAPs and effectors have membrane-targeting elements, which often contribute to regulating the activity of the GTPase. Together, the canonical regulators of Rho GTPases thus create a complex network of interactions that determine the precise spatiotemporal activation of Rho GTPases, all of which can potentially contribute to impairing signaling in AID. However, this has remained poorly understood.

Here, we report the identification of a R186C mutation in the C-terminal region of Cdc42 in a young adult patient presenting an autoinflammatory phenotype who has developed a severe form of generalized pustular psoriasis following bone marrow transplant in childhood. We provide the molecular mechanisms demonstrating the pathogenicity of the R186C mutation which causes palmitoylation of the carboxy terminal domain responsible for Cdc42 trapping in the Golgi apparatus. We also link the systemic inflammatory phenotype of the patient to proinflammatory cytokines overproduction due to increased NF-κB activation. Altogether, these data indicate that mutations located in the C-terminal part of Cdc42 unbalance its shuttling between the Golgi and the plasma membrane. As a consequence, downstream signaling pathways depending on proper subcellular localization, such as actin polymerization and NF-κB signaling are impaired. Thus, our results definitively link mutations in the *CDC42* gene to a complex immune-hemato-autoinflammatory phenotype in humans.

## Results

### Clinical and genetic features

A.S. was born at term to healthy non-consanguineous Caucasian parents. He has developed, since the age of 12 days of life, episodes of fever, hepato-splenomegaly, pancytopenia, diffuse maculopapulous rash with many relapses and a progressive worsening of the general condition. At 2 months of life a new episode of pancytopenia led to several investigations showing an extra medullary hematopoiesis at puncture liver biopsy, a non-specific lymphoid hyperplasia at a lymph node’s biopsy and a poor bone marrow. The skin biopsy was non-specific with a moderate non-specific inflammatory infiltrate. A treatment with high doses of systemic steroids was introduced, with improvement but a corticoid-dependence. At 11 months, he displayed relapses of scaly erythematous skin lesions, fluctuating hepato-splenomegaly with cytolysis, severe impact on growth (−4DS). At 16 months, the patient’s health worsened with a moderate bone marrow fibrosis, persistence of skin flare-ups, episodes of fever and cytolysis with a non-specific inflammatory syndrome, the absence of auto-immunity, the absence of infections. A splenectomy was performed at 23 months, showing many outbreaks of hematopoiesis, with no evidence of a tumor pathology. An allogeneic bone marrow transplant was decided at 24 months, with a good immune reconstitution, a chimerism 100% donor. However, scaly cutaneous lesions appeared, as early as 6 days after the transplant, with an aspect of psoriasiform lesion at skin biopsy and the absence of arguments for a GVH disease. These lesions gradually worsened until an involvement of the entire body (erythroderma) at M8 of the transplant, and episodes of amicrobial pustular lesions with, at skin biopsy, aspects consistent with pustular psoriasis-type lesions (**Fig. 1A**). On the other hand, there was a total regression of fever and pancytopenia, while hepatomegaly and liver cytolysis persisted. The patient is now a young adult of 23 years old. He evolves to this day with a picture of permanent severe psoriasiform erythrodema (**Fig. 1B**), with flares of more pronounced painful skin inflammation (**Fig. 1C**), chronic cutaneous bacterial infection by staphylococcus aureus, chronic episodes of cytolysis. Viral infections and staphylococcal cutaneous infections worsen the organ inflammation. Clinically, aortic deficiency, mild dysmorphia. acquired hypophosphatemic rickets, growth retardation associated, a non-specific inflammatory syndrome, a variable monocytosis (>1000/mm^3^) hypereosinophilia (>600/mm^3^) and hyper IgE (>700) are observed. The clinical characteristics of the patient with flares of inflammation in different organs with episodes of fever and non-specific inflammatory syndromes, are in favor of an autoinflammatory disease. The various therapeutic trials with immunosuppressive treatments and/or biotherapies including anti-IL1 and anti TNF, tested in severe and pustular psoriasis, are in check after transient improvement (**Fig. 1D-E**).

**FIG 1.**
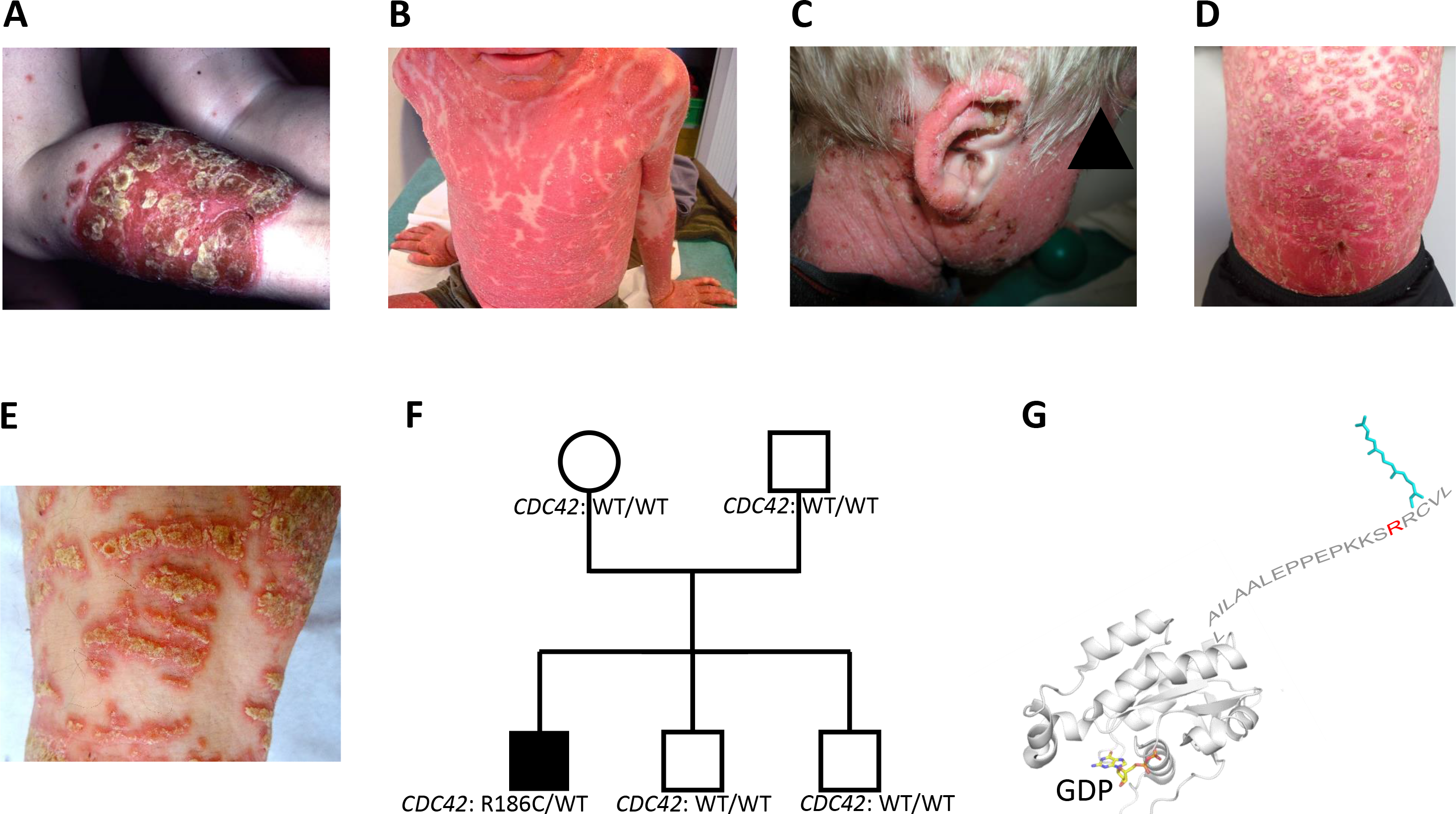
**A**, Pustular psoriasiform lesions in plaques, with a sudden onset after a viral episode, at the age of 4. Severe dermatitis (psoriasiform erythroderma) (**B**) with chronic crusted lesions and impetiginisation (**C**). Mild improvement of the skin with classical treatments for severe psoriasis (**D**), and chronic relapses of pustular psoriasiform lesions, with chronic flares of severe skin inflammation (**E**). **F**, Pedigree of the patient showing a *de novo* mono-allelic mutation in *CDC42*. The affected male subject is indicated as a black square. His mother is shown as an open circle. His father and two brothers are symbolized by open squares. *WT,* Wild type. **G**, Structure of human Cdc42 showing the R186 (in red) amino acid in the C-terminal hypervariable region (HVR). This region is modified by a geranyl-geranyl lipid (in cyan) attached to C188, which is critical for the attachment of GDP-bound Cdc42 to the membrane. The GTPase core is shown in cartoon representation. The hypervariable region is depicted by its sequence.

We performed exomes with DNA from the patient and his parents (**Sup Fig. 1A**). As the patient has been transplanted, we used DNA extracted from his fibroblasts in which we checked the different genetic background from the donor one. We first selected heterozygous neo-variations carried only by the patient and absent in parents and absent from public databases (dbSNP, 1000 genoms and ExAC). After segregation in the whole family including the two healthy brothers, we retained the Chr1:22417990C>T mutation in the *CDC42* gene which changes an arginine into cysteine in position 186, just before the CAAX sequence. The mutation was absent in the parents and in the two healthy brothers (**Fig. 1F**, **Sup Fig. 1A**). The mutation affects only the ubiquitously expressed Cdc42 form (**Sup Fig. 1C**) because the brain form differs for the last ten amino acids due to alternative splicing ^21,22^ (**Sup Fig. 1D)**.

At the structural level, R186 is located in the hypervariable C-terminal region (HVR) which carries post-translational lipid anchors (**Fig. 1G**), and it contributes to electrostatic interactions with negatively charged phospholipids ^23^. Thus, the mutation may affect aspects of Cdc42 association to / dissociation from membranes.

### The mutant Cdc42 R186C protein is trapped in the Golgi apparatus

We first aimed at deciphering the impact of the R186C mutation on the subcellular localization of Cdc42. Thus, we expressed myc- or GFP-tagged wild-type (WT) or R186C Cdc42 in both the CEM cell line derived from the T cell lineage or the HBMEC adhesive endothelial cells derived from the bone marrow. The mutant protein accumulates in a restricted cytoplasmic region, unlike the WT protein that exhibits a diffuse intracellular localization (**Fig. 2A**). Using the GM130 Golgi marker, we show that Cdc42 R186C is highly co-localized with the Golgi apparatus, contrary to the WT protein (**Fig. 2B-C** left). The quantification of the co-localization degree between the Golgi apparatus and Cdc42 was performed using the Pearson’s coefficient (PC). Our results show that cells transfected with WT protein have an average PC of 0.5 while it is significantly higher for the mutant protein (0.8-0.9) (**Fig. 2B-C** right, **Fig. 2D**) indicating that the mutation causes an accumulation of the protein in the Golgi.

**FIG 2.**
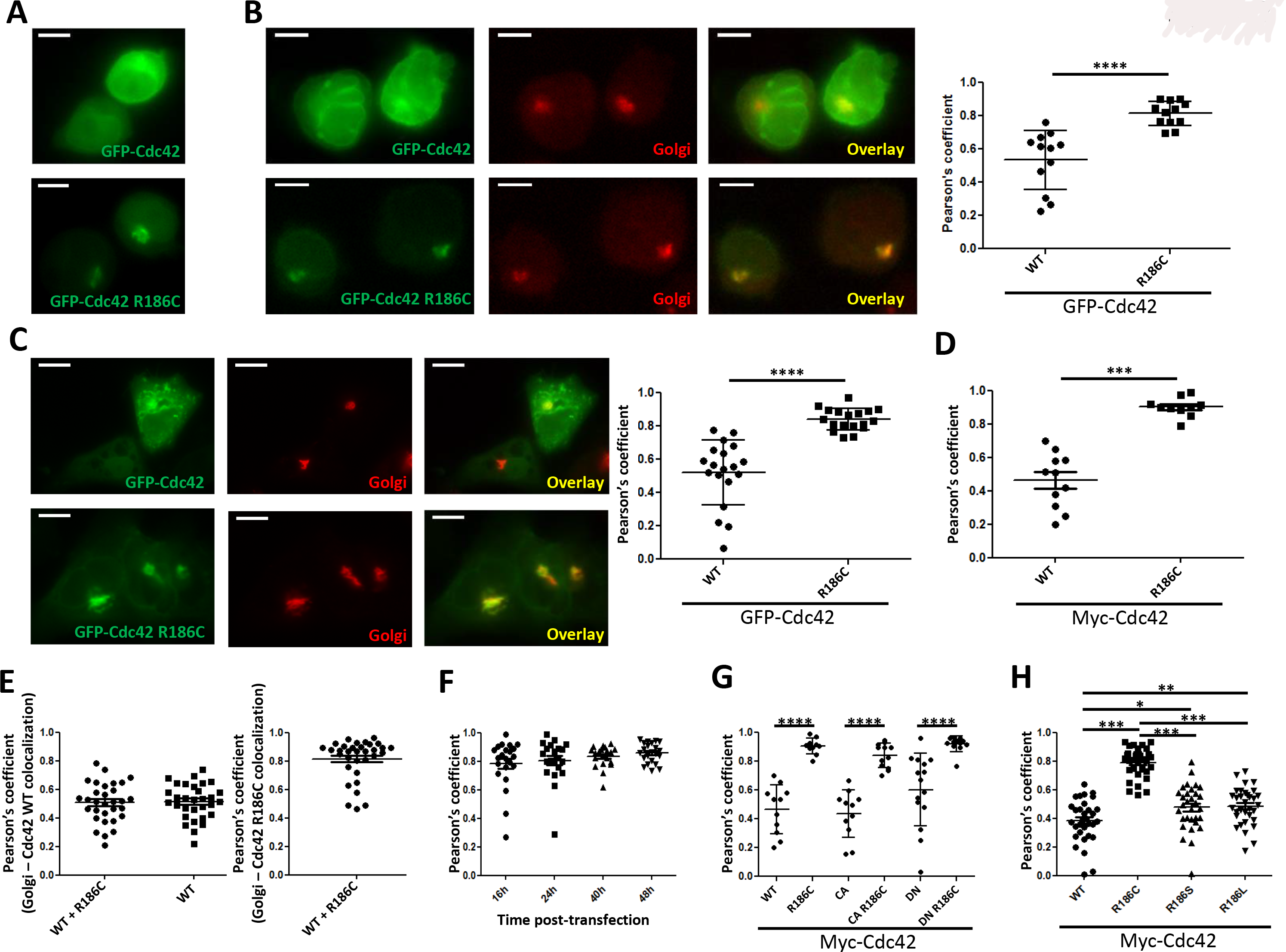
**A**, Subcellular localization of GFP-Cdc42 (top) or GFP-Cdc42 R186C (bottom) in live CEM cells. Staining of the Golgi resident GM130 protein (red) in CEM (**B**) and HBMEC (**C**) cells expressing GFP-Cdc42 (top) or GFP-Cdc42 R186C (bottom). The overlay of green and red fluorescence is also shown. The graphs on the right represent the Pearson’s coefficient measured in each condition. Scale bars are 5 μm (CEM) or 20 μm (HBMEC). **D**, The Pearson’s coefficient was also quantified on CEM cells expressing Myc-Cdc42 or Myc-Cdc42 R186C. **E**, HBMEC cells expressing either Myc-Cdc42 WT and GFP-Cdc42 R186C or Myc-Cdc42 WT alone were analyzed by microscopy to determine the degree of Golgi co-localization with WT (left) or R186C (right) Cdc42. **F**, Degree of co-localization between the Golgi apparatus and Myc-Cdc42 R186C analyzed between 16 and 48 hours after transfection. **G**, Measurement of the Pearson’s coefficient for constitutively active (CA) or dominant negative (DN) Cdc42 forms associated or not with the R186C substitution. **H**, Quantification of the degree of co-localisation between the Golgi and Myc-tagged WT or mutant Cdc42 in CEM cells. The graphs are representative of at least three independent experiments. *: p<0.05, **: p<0.01, ***: p<0.001, ****: p<0.0001.

Because, the Cdc42 mutation observed in this patient is heterozygous, we studied whether the mutant protein could influence the localization of the WT protein. For this purpose, both Cdc42 forms were co-transfected in HBMEC cells. The obtained PC values indicates that both WT and mutant Cdc42 localize in cells independently of each other (**Fig. 2E**), further showing a strong and stable (**Fig. 2F**) Golgi localization of Cdc42 R186C.

We then wanted to determine whether the abnormal subcellular localization of Cdc42 R186C depended on its state of activation. For this purpose, we used plasmids bearing both the R186C mutation and either the constitutively active (CA) Q61L mutation which mimics GTP-loaded Cdc42 or the dominant negative (DN) T17N mutation mimicking the inactive GDP-bound Cdc42 form. No difference was observed between active or inactive Cdc42 (**Fig. 2G**). These data indicate that the Golgi retention of Cdc42 R186C is independent of its activity status.

We asked whether Golgi retention of the mutant was solely due to removal of a positively charged residue or was specific of the mutation to cysteine. We thus designed R186S and R186L mutants. Serine has a structure close to a cysteine but it cannot be modified by lipid addition and Leucine has a hydrophobic side chain. Remarkably, R186S and R186L Cdc42 exhibit a low level of Golgi co-localization as shown by means PC of 0.5 in both cases, only slightly higher than that to WT Cdc42 (**Fig. 2H**). The same general observations were drawn from experiments performed in HBMEC cells (**Sup Fig. 2A**) and in resting primary human T cells (**Sup Fig. 2B**). All these results indicate that it is the specific mutation to cysteine in position 186 that is responsible for the presence of mutant Cdc42 in the Golgi apparatus.

### Cdc42 R186C accumulation in the Golgi is due to a palmitoylation

We next aimed at elucidating the molecular mechanisms responsible for Cdc42 R186C retention in the Golgi apparatus. Alignments between the C-terminal amino acid sequence of the Cdc42 R186C protein with those of other small G proteins ^24^, revealed that the R186C mutation introduces a cysteine in a position that is palmitoylated in the case of H-Ras (**Sup Fig. 3**).

Therefore, we next wondered whether cysteine 186 in mutant Cdc42 could also be palmitoylated. To directly test this hypothesis, we performed immunoprecipitation of Cdc42 from cells cultured in the presence of radioactive palmitate. The results demonstrated that Cdc42 R186C is palmitoylated whereas WT Cdc42 or the R186S mutant are not (**Fig. 3A**). This strongly suggests that the cysteine present in position 186 is palmitoylated.

**FIG 3.**
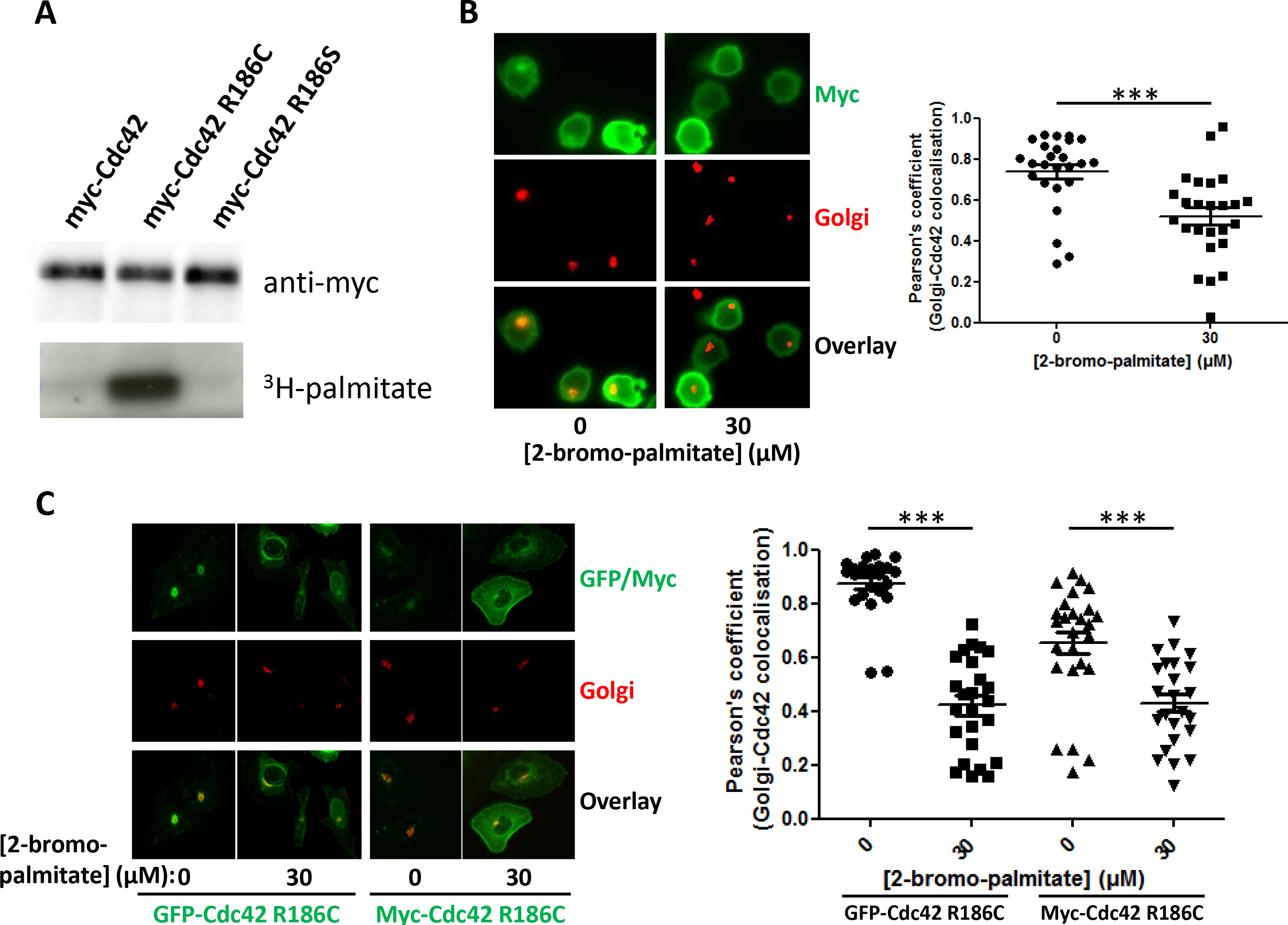
**A**, Western blot showing the amount of palmitate borne by Myc-tagged Cdc42, Cdc42 R186C or Cdc42 R186S. **B**, Localization of the Golgi apparatus and Myc-Cdc42 R186C in CEM cells treated or not with 30 μM of 2-bromo-palmitate. **C**, HBMEC cells expressing GFP (left) or Myc (right) -tagged Cdc42 R186C treated or not with 30 μM of 2-bromo-palmitate were analyzed in the same way. The quantifications of Pearson’s coefficients measured in each condition are shown on the right. Each graph is representative of three independent experiments. ***: p<0.001.

Next, we aimed at determining whether this aberrant palmitoylation could account for the Golgi accumulation of Cdc42 R186C. Using 2-bromo-palmitate, a palmitate analogue that inhibits palmitoyl transferases, we show that blocking palmitoylation activity induces a loss of Golgi localization of Cdc42 R186C, as shown by a decrease of the mean PC from 0.7 to 0.5 (**Fig. 3B**). HBMEC cells expressing GFP or Myc -tagged Cdc42 R186C also show a decrease in the PC from 0.9 to 0.4 for GFP-Cdc42 R186C and from 0.7 to 0.4 for Myc-Cdc42 R186C upon inhibition of palmitoylation (**Fig. 3C**).

Together, these results demonstrate that the cysteine residue in position 186 causes the palmitoylation of Cdc42 and that this prenylation is responsible for the retention of mutant Cdc42 in the Golgi apparatus.

### Cdc42 R186C fails to interact with GDI1

Because Cdc42 R186C displays a localization defect, we hypothesized that it may lose or gain molecular partners. Therefore, in order to identify the interactome of mutant Cdc42 in an unbiased fashion, we performed a mass spectrometry (MS) analysis to compare the identity of proteins able to bind WT *versus* R186C Cdc42.

The binding of effectors to the Cdc42 mutant varied highly between effectors. Some effectors, such as IQGAP3 bound equally well both WT and mutant Cdc42 (**Fig. 4A**). Others, such as the still poorly studied Cdc42EP and Cdc42SE families, bound the mutant better than the wild type protein. Conversely, partners encoded by the *FIGNL2*, *FAM89A*, *IQGAP1/2* and *MYL6* genes bound less to mutant Cdc42. Importantly, interaction with GDI1, which shuttles Cdc42 between the Golgi and the plasma membrane, was significantly reduced by the mutation. Because GDI proteins play a crucial role in the intracellular trafficking of Rho GTPases ^16,18,19,25^, we focused our analysis on the Cdc42-GDI interaction. MS analysis showed a statistically significant decrease in GDI1 binding of the mutant form of Cdc42 compared to the WT.

**FIG 4.**
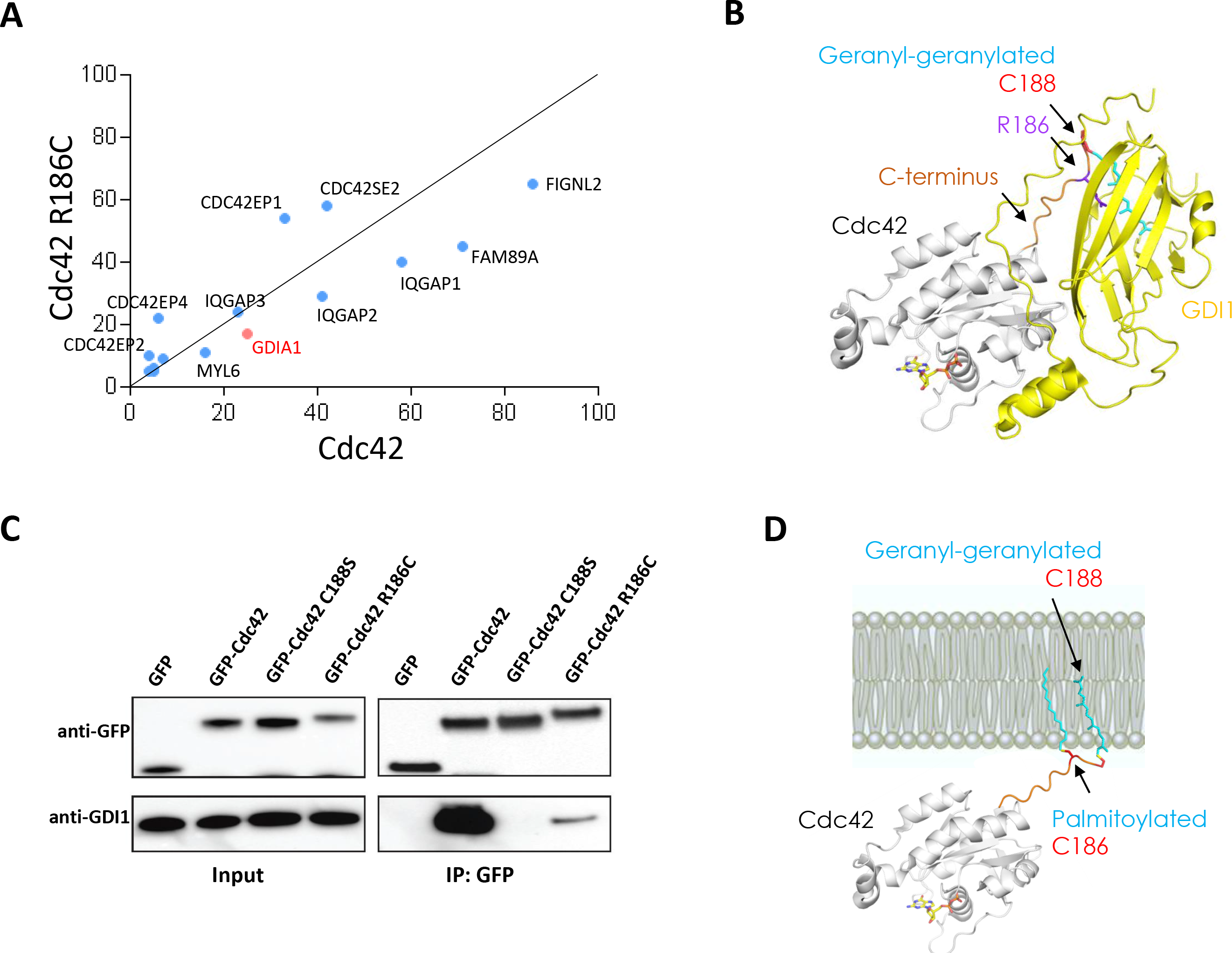
**A**, MS analysis shown as a correlation plot representing the level of abundance of identified interactors to WT or R186C Cdc42 normalized over GFP. All proteins have a peptide ratio ≥ 4 over GFP, a p-value ≤ 0.05 and peptides used to calculate significance are ≥ 6. **B**, Structure of the Cdc42-GDI1 complex. R186 (in purple) is buried in a groove of GDI1 (in yellow), which regulates the alternation of Cdc42 between the membrane and the cytosol by forming a soluble Cdc42-GDI complex. **C**, GFP (negative control) or GFP-tagged Cdc42 (WT, non geranyl-geranylated C188S mutant or R186C mutant) expressed in HEK cells were immuno-precipitated (IP) and tested for their ability to bind GDI1 by Western blot. The input fraction shows the amount of GFP, GFP-tagged Cdc42 proteins and GDI1 present in the different whole cell lysates. One representative experiment out of three is shown. **D**, The R186 to cysteine (C186) mutation allows its modification by a second lipid (palmitate). This second lipid cannot be accommodated by the R186 pocket of GDI1 and it enhances the interaction of Cdc42 R186C with the membrane, impairing the cytosol/membrane alternation.

The crystallographic structure of the Cdc42/GDI1 complex shows that GDI1 interacts extensively with the lipidated HVR of Cdc42 and buries the gerany-geranyl lipid in a hydrophobic pocket ^26^. Importantly, the R186 residue is also buried by GDI1 (**Fig. 4B**). Therefore, the substitution of Arg 186 by a palmitoylated cysteine at this particular position could interfere directly with GDI1 binding.

To address this issue, we biochemically compared the amount of GDI1 bound to GFP-tagged WT or R186C Cdc42. Negative controls were GFP alone and the C188S Cdc42 mutant which, by lacking geranyl-geranylation ^22^, cannot bind GDI1 (**Fig. 4C**). As reported, WT Cdc42 exhibits a strong binding to GDI1, and this was abolished by the control C188S mutation. Remarkably, the interaction of Cdc42 R186C with GDI1 was drastically reduced (**Fig. 4C**).

Together, our data suggest that the R186C mutation results in an increased interaction of the mutant with Golgi membranes arising from 1) aberrant palmitoylation that increases affinity for the membrane 2) decreased dissociation by GDIs due to removal of a major protein-protein interaction and 3) impaired interactions with negatively charged lipids that are abundant at the plasma membrane, which favors interactions with Golgi membranes (**Fig. 4D**).

### Cdc42 R186C is defective in actin polymerization

We next wondered what could be the functional consequences of the aberrant localization of Cdc42 R186C on its function.

Because Cdc42 classically controls actin filaments (F-actin) polymerization, we first asked whether actin polymerization was affected. We used flow cytometry to quantify F-actin levels in CEM cells expressing Myc-tagged WT *versus* R186C Cdc42. Both Myc^+^ WT and R186C cells exhibited a mean increase in F-actin levels of 35 % compared to cells that do not express these proteins (**Fig. 5A**).

**FIG 5.**
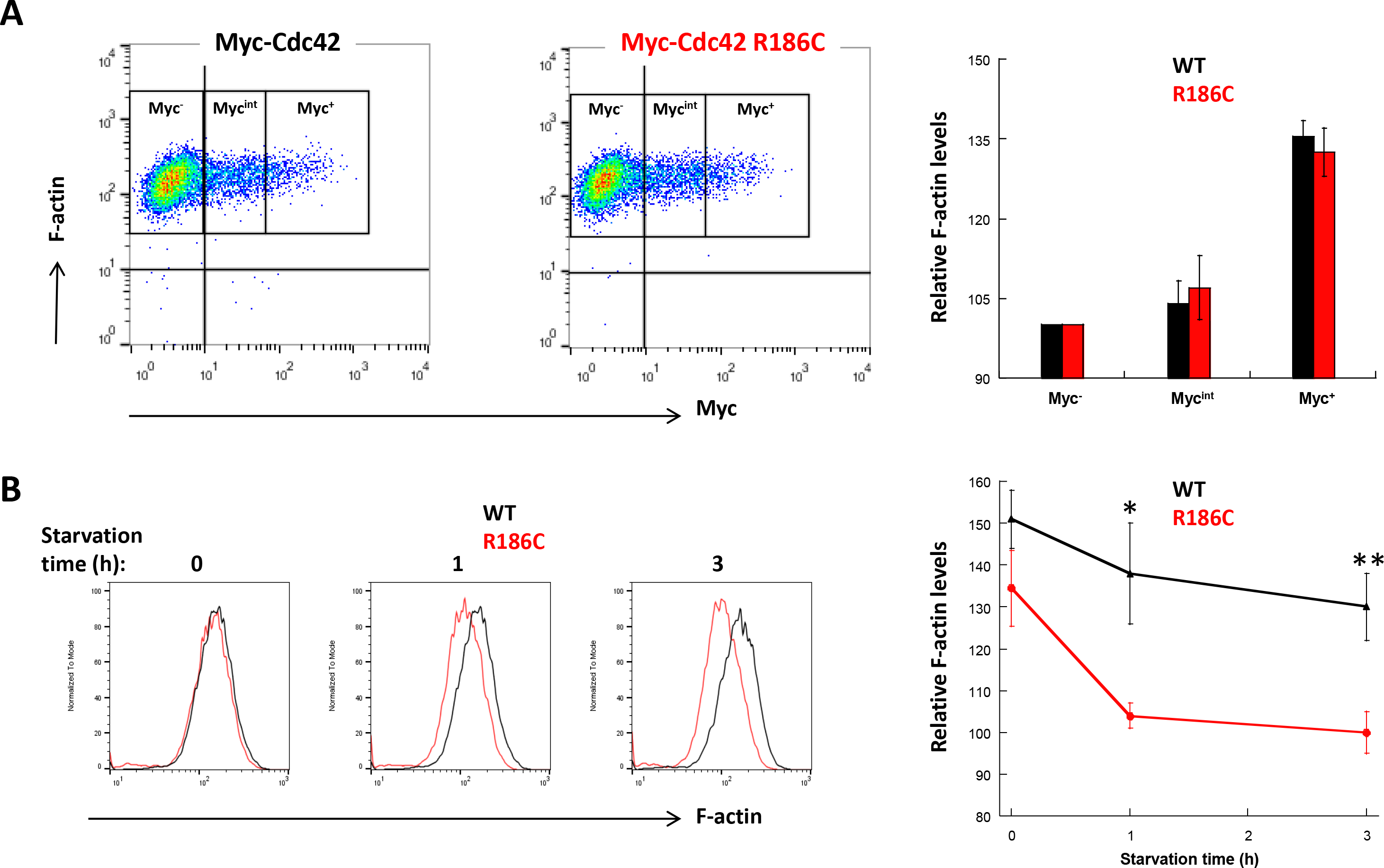
**A**, Dot plot showing the amount of F-actin as a function of myc expression in CEM cells transfected with Myc-Cdc42 (left) or Myc-Cdc42 R186C (right). Three regions were outlined depending on Myc expression: Myc^−^ cells do not overexpress Cdc42, Myc^int^ cells express intermediate levels of plasmid-encoded Cdc42 protein, and Myc^+^ cells express high levels of either Myc-Cdc42 or Myc-Cdc42 R186C. The graph on the right shows the means +/− SE of F-actin levels measured in five independent experiments after normalization to 100 for the Myc^−^ conditions. **B**, Flow cytometry histograms showing the F-actin levels measured in WT Myc-Cdc42^+^ (black) *versus* Myc-Cdc42 R186C^+^ (red) CEM cells after 1 or 3 hours of serum starvation. The graph on the right shows the means +/− SE of F-actin levels measured in Myc^+^ CEM cells from four independent experiments after normalization to 100 for the Myc^−^ conditions. *: p<0.05, **: p<0.01.

However, when cells were placed in serum starvation to decrease endogenous GEFs activation, Cdc42 R186C^+^ cells quickly failed to sustain actin polymerization compared to WT Cdc42^+^ cells (**Fig. 5B**). The drop in F-actin levels observed in cells expressing Cdc42 R186C could be observed as early as 1 hr upon serum starvation and was still observed at 3 hrs.

Thus, these results indicate that the R186C mutation reduces the ability of Cdc42 to induce F-actin polymerization and confirm that Cdc42 mutation impairs the protein ability to activate signaling cascade at the plasma membrane, where it promotes actin polymerization.

### Cdc42 R186C confers a pro-inflammatory phenotype

In order to relate our observations to the systemic inflammatory phenotype of the patient, we assessed the production of proinflammatory cytokines by patient fibroblasts in comparison to independent healthy individuals upon several ligand stimulation. Patient fibroblasts overproduced significantly the 10 proinflammatory cytokines compared to controls (**Fig. 6A**). No variation was observed for the anti-inflammatory cytokine IL-13 as well as for IL-17A (**Sup Fig 4**). Because a cross-talk between the NF-κB and Rho signaling pathways has been previously reported, we also used a specific anti-NF-κB inhibitor that impairs IKKβ/IKKχ binding required for IKK kinase activity during cell stimulation. Interestingly, we show that this inhibitor restored the cell inflammatory phenotype (**Fig. 6B**) indicating that overproduction of proinflammatory cytokines in the patient fibroblasts is NF-κB -dependent. We next measured NF-κB nuclear translocation in control or patient fibroblasts stimulated by LPS. We observed that patient cells exhibited an almost two-fold increase in p65 NF-κB nuclear translocation compared to control cells (**Fig. 6C**). Because NF-κB activity is principally mediated by a set of phosphorylation events, we thus assessed phosphorylation of p65 at serine 529 in control and patient fibroblasts stimulated with TNFα. We show an increased p65 phosphorylation in non-stimulated patient fibroblasts over control ones which is more important after TNFα treatment (**Fig. 6D**). Finally, in order to link Cdc42 to NF-κB activation, we performed a NF-κB transactivation assay into HEK 293T cells expressing inactive, active and the R186C Cdc42 mutant. We observe that a constitutively active Cdc42 stimulates NF-κB whereas no stimulation was obtained with the dominant-negative Cdc42 form. Remarkably, Cdc42 carrying the R186C mutation activates NF-κB more than the wild type form (**Fig. 6E**).

**FIG 6.**
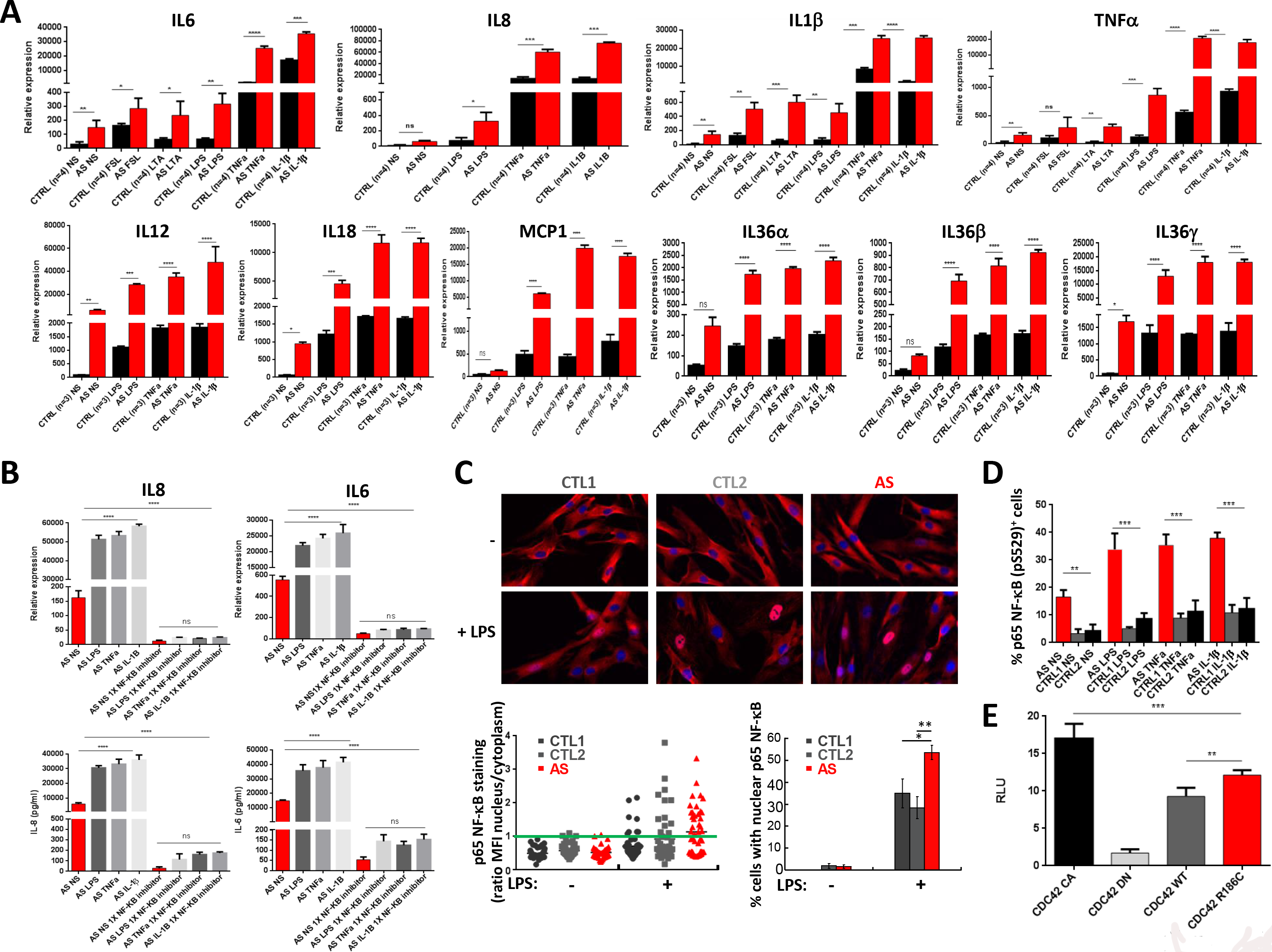
**A**, Relative expression of IL-6, IL-8, IL-1β, TNFα IL-12, IL-18, MCP-1, IL-36α, IL-36β and IL-36 transcripts in AS and healthy control fibroblasts unstimulated or stimulated with LPS, TNFα, IL-1β FSL-1 or LTA. **B**, qPCR (top) and ELISA (bottom) of IL6 and IL8 expression in unstimulated or stimulated control or AS fibroblasts in the absence or presence of NF-κB inhibitor. **C**, top: Images of control or patient fibroblasts stimulated or not with LPS and stained for p65 NF-κB (red) and Hoechst (blue). Translocated p65 NF-κB staining appears as a pink nucleus. Bottom left: Plot showing the nucleus to cytoplasm ratio of p65 NF-κB stainings from 47 to 53 randomly-imaged cells. Cells exhibiting a ratio above 1 (green line) were considered to be cells showing p65 NF-κB nuclear translocation. Bottom right: Mean +/− SE of the percentage of cells showing p65 NF-κB in the nucleus from five independent experiments. **D**, Phospho-p65 (p65 PS529) detection by flow cytometry in patient fibroblasts upon 30 min of TNFα stimulation compared to two healthy individuals. Means +/− SE from three independent experiments. **E**, NF-κB luciferase activity in cells transfected with different forms of Cdc42. *: p<0.05, **: p<0.01, ***: p<0.001 and ****: p<0.0001.

## Discussion

Here, we describe the case of a patient with a severe autoinflammatory syndrome carrying a R186C mutation in the ubiquitously expressed Cdc42 isoform. The main clinical characteristics of the patient are a severe dermatitis since birth, with initially flares of a non-specific urticarial and scaling rash, then well delimited psoriasiform plaque and finally chronic psoriasiform erythroderma resistant to the different lines of treatment, flares of hepatomegaly and cytolysis, mild dysmorphia, and a permanent non-specific inflammatory syndrome with variable monocytosis, absence of auto-immunity and, at adult period, a mild hyper eosinophilia and hyper IgE that could be related to a skin barrier impairment. No major infections nor allergies are observed, but a chronic staphylococcal colonization acts as a triggering factor for worsening skin inflammation.

Our results show that the mutation of R186 by cysteine in the HDR, a region which is increasingly recognized for small GTPase localization and functional specificity ^20,27^, renders it permissive to palmitoylation, resulting in a dually prenylated mutant. This observation fully explains the massive displacement of Cdc42 from the cytosol to the Golgi and suggests how it alters Cdc42 signaling.

In wild-type Cdc42, R186 is a critical determinant for the regulated membrane/cytosol switch by forming dual interactions with either the membrane in its active, membrane-associated form ^23^ or GDI in its inactive, cytosolic form ^26^. Accordingly, replacement of R186 by cysteine and its posttranslational palmitoylation is expected to induce multiple defects in the functional cytosol/membrane alternation of Cdc42. First, palmitoylated C186 decreases the interaction with GDI1, hence impairs extraction of the mutant from the membrane resulting in a more stable interaction with membranes. Second, replacement of R186 by palmitoylated C186 should affect the membrane preference of Cdc42, both by removing a positively charged residue that favors the recognition of negatively charged phosphatidylinositides that are enriched at the plasma membrane ^23^ and by adding a lipid known to support trafficking through the Golgi (for example: reviewed in ^28^). Thus, both contributions are likely to determine why the Cdc42 mutant appears mostly trapped at the Golgi.

Our data using constitutively active and inactive forms of the R186C mutant indicate that localization of the mutant at the Golgi is largely independent from its nucleotide status. This is consistent with the fact that the mutation is remote from the GDP/GTP-binding site and should thus not affect directly the ability of the mutant to bind to GEFs, GAPs and effectors *in vitro*. However, it is striking that the mutation affects binding to effectors in a differential manner in the cell. A plausible explanation is that abnormal Golgi membrane targeting of the mutant concurrently upregulates its interactions with Golgi-localized effectors and downregulates those with plasma membrane localization. Accordingly, it suggests that physiological effects of the mutation could result from an imbalance between Cdc42 signaling pathways that normally work in concert on different endomembanes in the cell. It is interesting that the brain-specific Cdc42 isoform, which differs from the ubiquitous Cdc42 mostly by the sequence of its HVR, has features in common with the C186 mutant, including a dual lipidation and a marked vesicular distribution that contrasts with the mostly cytoplasmic distribution of ubiquitous Cdc42 ^29^. Thus, the distinct localization pattern of brain Cdc42 may endow it with a still different set of effectors through colocalization, hence function through alternative signaling pathways.

The present study that provides in depth molecular understanding of the Cdc42 R186C mutation further supports recent work that described four additional patients with autoinflammatory syndromes presenting C-terminal variants of Cdc42 ^11^: one mutation causes read-through of the stop codon and adds 24 amino acids, the same R186C mutation and finally, a C188Y mutation of the CAAX sequence. Interestingly, the C188S mutation produces a non geranyl-geranylated soluble form of Cdc42 that does not interact with GDI1. Consequently, the C188Y version of Cdc42 is most likely free to diffuse in the cytosol too. Interestingly, the dually lipidated R186C and non-lipidated C188Y are both likely to present a defect in plasma membrane anchoring. Our results showing that Cdc42 R186C fails to sustain actin polymerization in serum-starved cells are in agreement with this idea of mutant Cdc42 being less easily activated. In addition, the actin polymerization defect observed is compatible with the importance of actin dynamics in the activation of the inflammasome ^30^. Moreover, it has been demonstrated *via* data generated in both mouse and human that autoinflammation could be caused by mutations in the actin-regulatory gene *WDR1* ^31^.

A question remains: How all these C terminal Cdc42 mutations that preclude plasma membrane anchoring elicit an exacerbated inflammatory phenotype? Interestingly, a similar situation has been reported for RhoA: a defect in RhoA geranyl-geranylation has been shown to inhibit RhoA activity, activate the pyrin inflammasome and induce high levels of IL-1β production ^6–8^. In our study, the pro-inflammatory phenotype observed in our patient led us to investigate the NF-kB signaling pathway. It has been shown that Cdc42 signaling acts as a mediator of chronic inflammation associated with endothelial senescence, a feature of the inflammatory atherosclerosis disease ^32^. In these cells, inhibition of Cdc42 *via* NF-κB signaling attenuated the sustained up-regulation of pro-inflammatory genes. NF-κB signaling pathways have also been involved in a human inherited disorder characterized by chronic autoinflammation, and immunodeficiency related to loss-of-function mutations in HOIL-1 ^33^, a component of the linear ubiquitination chain assembly complex (LUBAC). Defects in A20 and Otulin also impair NF-κB ubiquitination and give rise to autoinflammatory diseases ^34^. In our patient, the results strongly support a direct link between Cdc42 and NF-κB. We indeed showed that TNFα-mediated NF-κB activity appears to be dependent on a functional Rho GTPases network, and probably on Cdc42 itself, as TNFα was unable to activate NF-κB in the presence of a dominant negative form of Cdc42 (data not shown). The extreme severity of the psoriasiform dermatitis exhibited by our patient is reminiscent of what is observed in mouse models of NF-κB invalidation in basal keratinocytes ^35,36^. For instance, a hypothesis in the *nemo* skin conditional knock-out mouse ^37^ is the interplay between mutated keratinocytes with wild type immune cells. A similar phenomenon could be suggested in our allogeneic bone marrow transplanted patient whose immune cells are normal in an otherwise mutated cell background. Further studies are needed to integrate various consequences of Cdc42 mutants which probably arise in a large spectrum of clinical phenotypes.

## Materials and methods

### Constructs

The pEGFP-C3 vector was from Clontech. The pEGFP-C3-Cdc42 plasmid was provided by L.I. Salazar-Fontana ^38^. The pRK5-myc-Cdc42 plasmid and the constitutively active L61Q and dominant negative T17N variants were all obtained from Addgene. The Cdc42 mutants (R186C, R186S, R186L or C188S) were generated by site-directed mutagenesis (Quickchange kit, Agilent Technologies).

### Cells

The lymphoblastoid T cell line CEM was grown in RPMI 1640 + Glutamax medium (Gibco) supplemented with 10 % heat-inactivated fetal calf serum (FCS), antibiotics (50 U/ml penicillin and 50 μg/ml streptomycin from Gibco), 10 mM sodium pyruvate (Gibco) and 10 mM hepes (Gibco).

Primary human peripheral blood T-cells (PBT) were purified from the blood of healthy donors provided by *Etablissement Français du Sang* using a Ficoll gradient separation before negative selection with a cocktail of antibodies (EasySep Human T cell isolation kit, Stem Cell) according to the manufacturer’s recommendation. PBT were grown in a complete RPMI 1640 supplemented by 10 % human serum AB.

Primary human fibroblasts from the patient carrying the Cdc42 R186C mutation were obtained by a biopsy of the skin. Control fibroblasts from healthy donors were similarly obtained.

HBMEC, HEK, RPE1 cells and primary human fibroblasts were cultivated in complete DMEM medium (Gibco).

### Transfections

2.10^6^ CEM cells were centrifuged for 5 minutes at 1200 rpm and washed in PBS 1X (Gibco). They were then transfected by nucleofection with 5 μg DNA in 100 μl of Cell Line Nucleofector Solution V (Lonza) using the C-016 program (Amaxa Biosystems). After transfection, 500 μl of complete RPMI medium was added to the cells, which were then deposited in 6-well plates containing 2 ml of medium. The plate was incubated at least overnight.

The transfection of the HBMEC cells was carried out according to the same protocol but with 3 μg DNA for 0.5×10^6^ cells in 100 μl cells of Cell Line Nucleofector Solution V (Lonza) with the program U-015. For the co-transfection experiments, 0.5×10^6^ HBMEC cells were transfected with 2.5 μg of each of the plasmids of interest encoding for GFP-Cdc42 R186C together with either Myc-Cdc42 WT or an empty vector.

PBT cells were transfected with 5 μg DNA for 5.10^6^ cells in 100 μl of Human T Cell Nucleofector Solution (Lonza) with the program U-014 (Amaxa Biosystems).

HEK cells were transfected with plasmids encoding GFP-tagged Cdc42 variants using Calcium Phosphate transfection or Lipofectamine LTX Kit (Life Technologies).

RPE1 cells were transfected using Fugene (Promega).

### Biochemistry

To detect palmitoylation, RPE1 cells transfected 48 hours earlier with different Myc-tagged constructs were starved for 1 hour at 37 †C in IM (Glasgow minimal essential medium buffered with 10 mM Hepes, pH 7.4), incubated for 2 hours at 37 †C in IM with 200 μCi/ml^3^H palmitic acid (9,10-^3^H(N)) (American Radiolabeled Chemicals, Inc.). The cells were washed, incubated in DMEM complete medium for 10 minutes, washed at 4†C three times with cold PBS and directly lysed for 30 min at 4 †C in Lysis Buffer [0.5% Nonidet P-40, 500 mM Tris pH 7.4, 20 mM EDTA, 10 mM NaF, 2 mM benzamidine and protease inhibitor cocktail (Roche)], and centrifuged for 3 min at 5000 rpm. Supernatants were subjected to preclearing with G sepharose beads prior immunoprecipitation reaction using an overnight incubation with anti-Myc affinity gel (Thermo Scientific). After immunoprecipitation, washed beads were incubated for 5 min at 90 †C in reducing sample buffer prior to 4-20 % gradient SDS-PAGE and revelation with a mouse anti-Myc 9E10 antibody (Covance). After SDS-PAGE, the gel was incubated in a fixative solution (25 % isopropanol, 65 % H2O, 10 % acetic acid), followed by a 30 min incubation with signal enhancer Amplify NAMP100 (GE Healthcare). The radiolabeled products were revealed using Typhoon phosphoimager.

For GDI1 co-immunoprecipitation assay, HEK cells were lysed using 50 mM TRIS base, Triton 2 %, 200 mM NaCl as well as 1 tablet/10 ml protease inhibitor Mini-complete, EDTA-free (Roche). After removal of insoluble fragments *via* centrifugation at 12,000 g for 25 min, lysates were incubated with 15 μl of GFP-Trap Agarose beads from Chromotek for 1 h at 4 †C on a rotary wheel. Beads were then washed three times for 10 min with wash buffer (250 mM NaCl and 0.1% Triton in PBS) followed by detection of GDI1 interaction with GFP-tagged Cdc42 variants using Western blot. Primary antibodies used were anti-GFP-HRP (Novus Biologicals) and anti-Rho GDIα (Santa Cruz Biotechnology).

### Mass Spectrometry

GFP immunoprecipitation assays were carried out as described above for GDI1 with one exception, here a special wash buffer was used to wash the Chromotek beads after binding assays. The mass spec wash buffer was prepared using 50 mM TRIS base, 150 mM NaCl, 1 mM EDTA and 2.5 mM MgCl_2_ and pH adjusted to 7.5. Following the final wash, beads were stored with wash buffer at 4 †C prior to depositing at the Institut Curie Mass Spectrometry and Proteomics facility (LSMP). For all the corresponding data analysis, the myProMS web server of the LSMP facility was used.

### Treatments of cells

CEM and HBMEC cells were incubated overnight in complete culture medium containing 30 μM of 2-bromo-palmitate (Sigma).

Cells were stimulated with FSL-1 (a TLR2/6 agonist; 1 μg/ml), LTA (a TLR 2 agonist; 10 μg/ml), LPS (a TLR4 agonist; 1 μg/ml), TNF-α (20 ng/ml) and IL-1β (20 ng/ml) for 24 hours.

### RNA extraction and cDNA preparation

Total RNA was extracted from patient and healthy control fibroblasts stimulated or left unstimulated with Qiagen RNA mini kit according to the manufacturer’s instructions. cDNA was synthesized thanks to iScript Ready to use cDNA Super mix (BioRad).

### Real time quantitative PCR

Real time quantitative PCR was performed with Syber Green PCR Master mix (Life Technologies) according to the manufacturer’s instructions. The results were read with the CFX384 Real Time System machine. Relative expression of mRNA was determined by the 2^−ΔΔC(t)^ method using *GAPDH* and *ACTIN* as housekeeping genes.

### The Enzyme Linked Immunosorbent Assay (ELISA)

Supernatants were collected and ELISA (InvivoGen) with different cytokines were performed according to manufacturer’s instructions. Absorbance results were read with the Infinite f200 pro TECAN machine.

### NF-κB luciferase activity

HEK 293 cells were transfected with NF-κB-dependent firefly luciferase vector, *Renilla* luciferase vector as an internal control and different constructions of Cdc42 following the manufacturer’s indications. The cells were lysed in passive lysis buffer and luciferase activity was read in the Infinite f200 pro TECAN machine.

### Immunofluorescence

After washing with PBS, cells were fixed with 4 % paraformaldehyde (PFA) (Electron Microscopy Sciences) for 10 minutes. They were then washed once in PBS containing 1 % Bovine Serum Albumin (BSA) (Sigma) and twice in a permeabilization buffer (PBS containing 0.1 % saponin (Fluka Biochemika) and 0.2 % BSA). Cells were incubated for 45 minutes with the following primary antibodies: anti-GM130 (Santa Cruz Biotechnology), anti-p65 NF-κB (Santa Cruz Biotechnology) or an anti-myc tag Alexa Fluor 488 antibody (Cell Signaling Technology; 9B11). After washing in the permeabilization buffer, cells were incubated for 30 minutes with secondary anti-goat or anti-mouse antibodies conjugated to Alexa Fluor 568 (Invitrogen). After an additional wash, cells were incubated with Hoechst (Sigma) for 5 min to stain nuclei.

For co-transfection experiments, the cells were stained according to the same protocol except for the myc staining that was performed using an unconjugated anti-myc tag antibody (Santa Cruz Biotechnology, 9E10) followed by an AMCA anti-mouse antibody (Jackson ImmunoResearch).

The images were acquired using a Nikon TE300 fluorescence microscope, a Cascade Photometrics camera and the Metamorph v7.8.9.0 software. Three oil-immersion objectives were used: 40×, 60× and 100×. Z-stack images were generated, and then, upon deconvolution and projection, the Pearson’s coefficient (PC) was measured on Fiji (ImageJ version 1.51u) using a macro containing the Coloc2 plugin. This coefficient measures the degree of overlap between two stainings and was used to quantify the degree of co-localization between Cdc42 and the Golgi apparatus. A PC equals 0 means that there is no co-localization between the two stainings. By contrast, a PC value of 1 means that there is a perfect co-localization between Cdc42 and the Golgi. The degree of p65 NF-κB nuclear translocation was quantified on Fiji from randomly acquired images by measuring the MFI of the staining in the nucleus divided by the MFI in the cytosol using same size regions. A ratio above 1 was considered to be the hallmark of a cell presenting p65 NF-κB nuclear translocation.

### Flow cytometry

Transfected CEM cells were used directly or serum starved during the indicated times in RPMI alone. Cells were then fixed and permeabilized as previously explained. Actin filaments were stained with 0.5 U/ml phalloidin Alexa Fluor 647 (Invitrogen). The amount of filamentous actin (F-actin) present in CEM cells was measured by flow cytometry (FACSCalibur, BD) and analyzed with the FlowjoV7 software.

Alexa Fluor 488 Mouse anti NF-κB p65 (pS529) from BD was used on fibroblasts.

### Statistical analysis

Statistical analyses were carried out using the graphPad Prism 5 software. The results represent the means +/− Standard Errors. The levels of significance were calculated by the Student t-test: *: p<0.05, **: p<0.01, ***: p<0.001, ****: p<0.0001.

## Acknowledgements

We thank Damarys Loew and Florent Dingli from the Laboratory of Proteomic Mass Spectrometry (LSMP) of Institut Curie.

This work was supported by Inserm, CNRS, Université Paris Descartes, Association pour la Recherche contre le Cancer (to JD), Ligue contre le Cancer (to SEM), Société Française de Dermatologie (to AS, JD and CB), Agence Nationale de la Recherche (RIDES to AS, JD and JC), the European Union Horizon 2020 Marie Skłodowska-Curie research and innovation programme (MSCA‐ITN‐2015‐675407 to SEM), the Institut Pasteur and the Fondation pour la Recherche Médicale (DEQ20150331694 to JC).

---

**Supplemental FIG 1.**
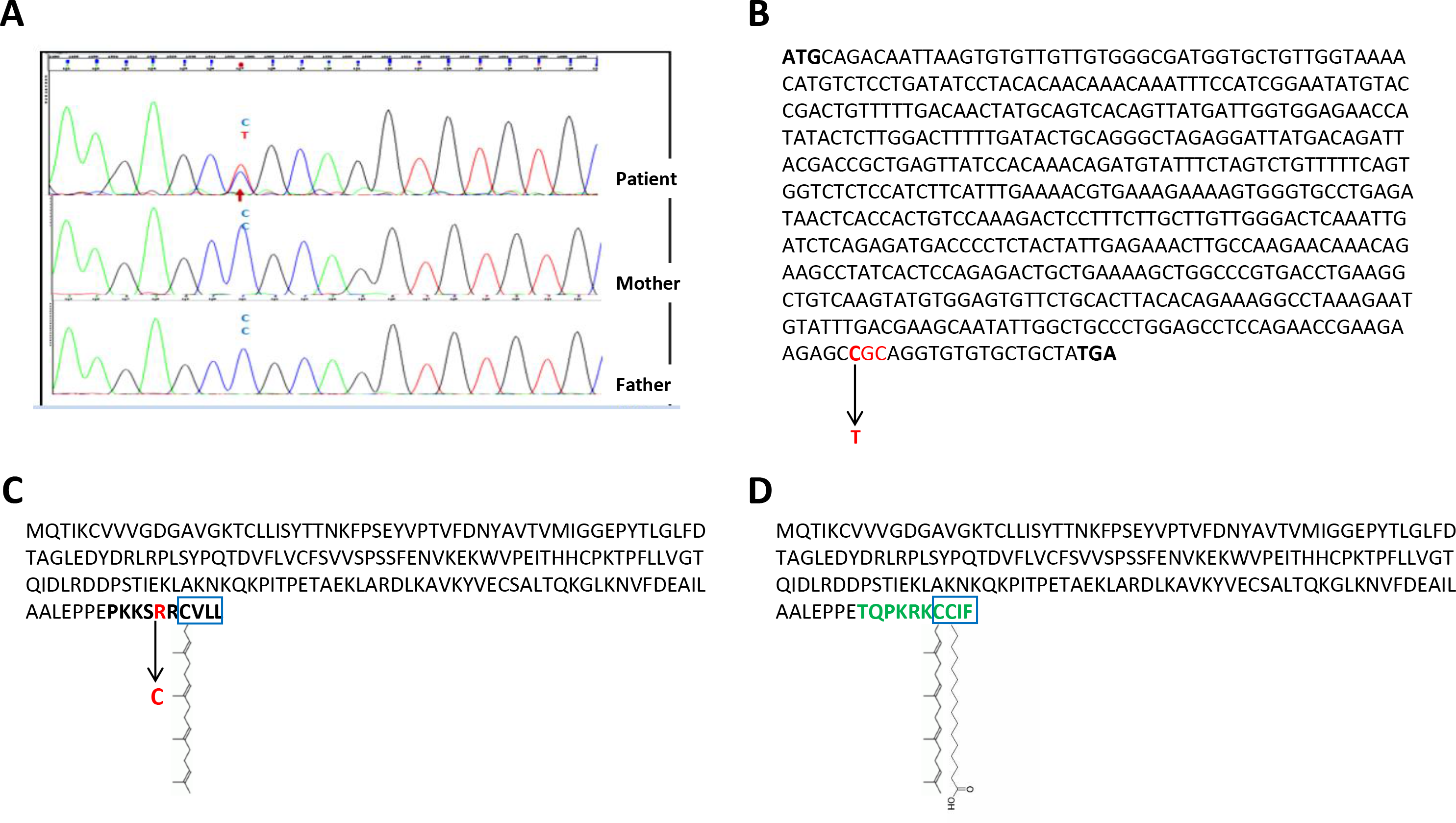
**A**, Electropherogram of the 3’ end coding sequence of the *CDC42* gene from the patient (top) and his two parents (bottom). Each peak corresponds to a nucleotide: adenine (A, green), cytosine (C, blue), thymine (T, red) and guanine (G, black). The red arrow indicates the mutation of a cytosine into a thymine on one allele. **B**, Nucleotides of the coding sequence of ubiquitous human Cdc42. The three nucleotides in red represent the codon modified by the C to T mutation. The start (ATG) and termination (TGA) codons are shown in bold character. **C**, Amino acid sequence of the ubiquitous human Cdc42 isoform. The blue box indicates the position of the CAAX sequence containing the geranyl-geranylation. The R186C mutation is shown in red. The last amino acids unique to this isoform are in bold character. **D**, Amino acid sequence of the brain-specific human Cdc42 isoform showing in green bold letters its unique C-terminal region. The blue box indicates the position of the CAAX sequence containing a geranyl-geranyl moiety on Cys188 and a palmitate on Cys189 ^22^.

**Supplemental FIG 2.**
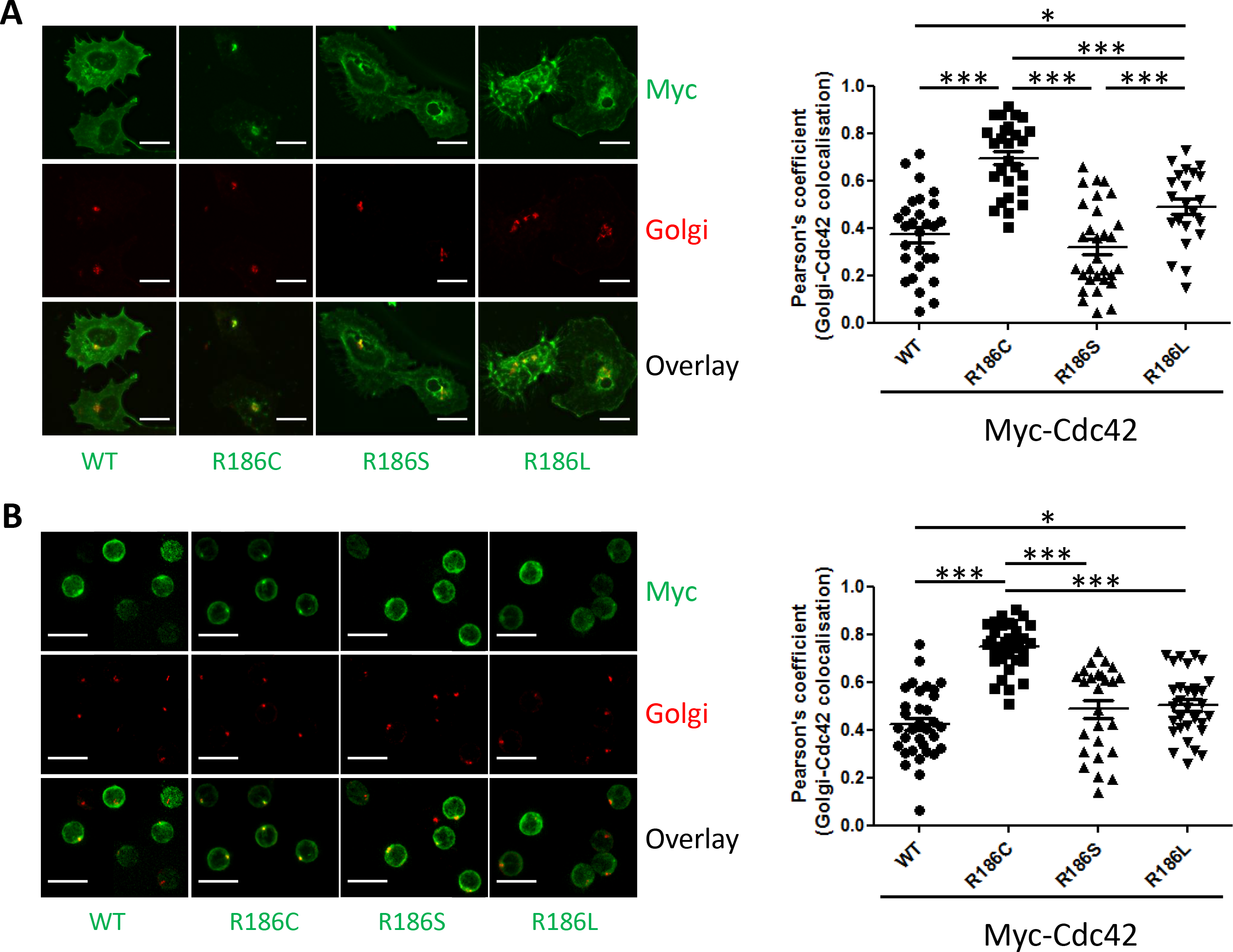
Left: Microscopy images of HBMEC (**A**) or primary resting human peripheral blood T lymphocytes (**B**) expressing Myc-tagged constructs encoding WT, R186C, R186S or R186L Cdc42. Stainings for Myc (green), Golgi-associated GM130 (red) and an overlay of both is shown for each condition. Scale bars: 20 μm (A) and 10 μm (B). Right: Quantification of the degree of co-localization between each Cdc42 forms and the Golgi apparatus is measured by the PC. *: p<0.05, ***: p<0.001.

**Supplemental FIG 3.**
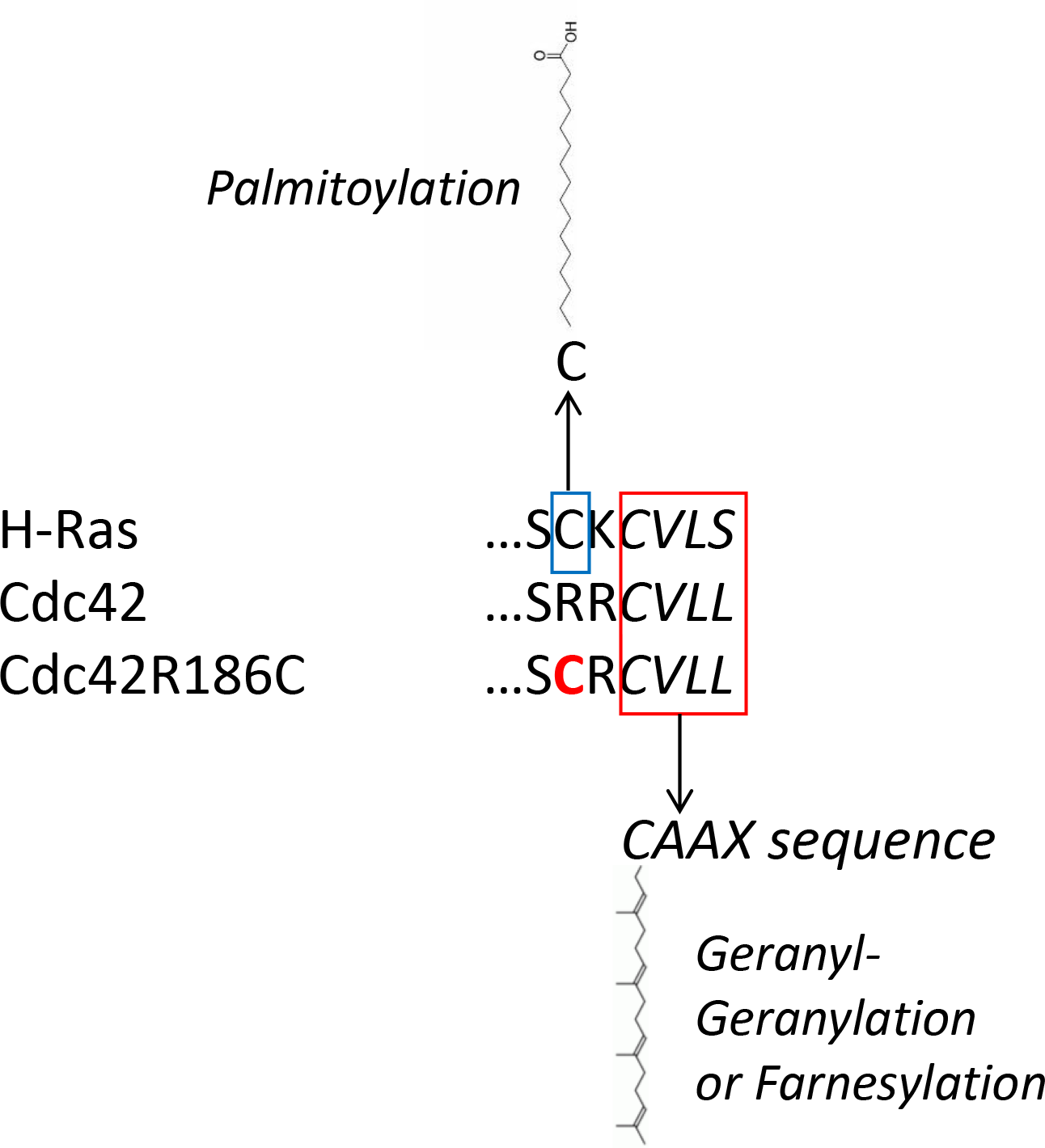
Alignment of C-terminal amino acid sequences of Cdc42, Cdc42 R186C and H-Ras. The Cyst residues in the CAAX sequences shown in a red box are geranyl-geranylated for Cdc42 or farnesylated for H-Ras. The H-Ras Cyst highlighted in a blue box is palmitoylated. Adapted from ^24^.

**FIG. S4.**
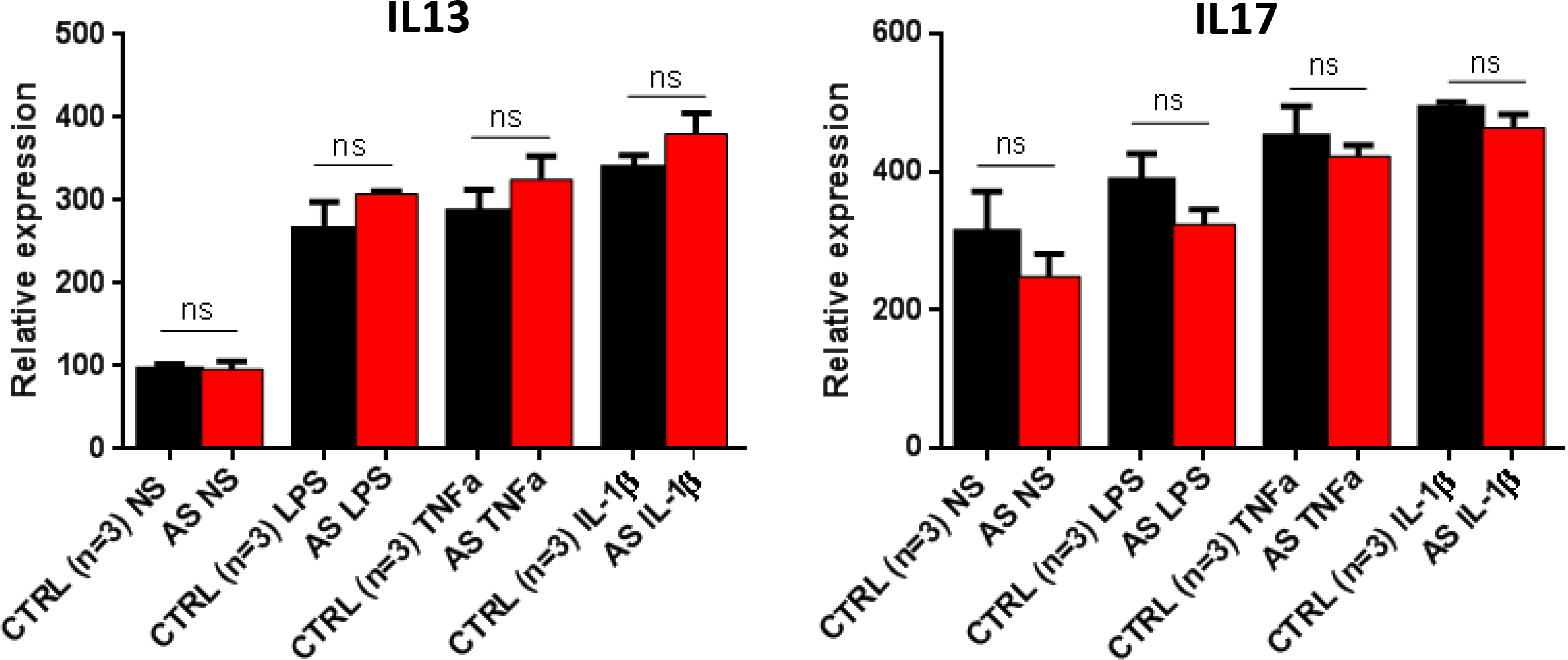
Relative expression of IL-13 and IL-17 transcripts in AS and healthy control fibroblasts unstimulated or stimulated with LPS, TNFα or IL-1β. ns: not significant.

